# A simple method to co-purify genomic DNA, RNA, and proteins for functional studies

**DOI:** 10.1101/467928

**Authors:** Jian Jiang, Junfei Ma, Bin Liu, Ying Wang

## Abstract

Understanding the regulation of gene expression, from the epigenetic modifications on genomes to posttranscriptional and translational controls, are critical for elucidating molecular mechanisms underlying distinct phenotypes in biology. With the rapid development of Multi-Omics analyses, it is desirable to minimize sample variations by using DNA, RNA, and proteins co-purified from the same samples. Currently, most of the co-purification protocols rely on Tri Reagent (Trizol as a common representative) and require protein precipitation and dissolving steps, which render difficulties in experimental handling and high-throughput analyses. Here, we established a simple and robust method to minimize the precipitation steps and yield ready-to-use RNA and protein in solutions. This method can be applied to samples in small quantity, such as protoplasts. We demonstrated that the protoplast system equipped with this method may facilitate studies on viroid biogenesis. Given the ease and the robustness of this new method, it will have broad applications for plant research and other disciplines in molecular biology.

## Introduction

The flow of genetic information from genome to transcriptome and then proteins dictates the phenotype in all organisms (Crick, 1970). The genomic *cis*-regulatory elements and epigenetic hallmarks (e.g., cytosine methylation) in genomic DNA sequences may help elucidate transcriptional regulations of genes expression, the effects of which may impact the corresponding mRNA and protein levels (Hollick, 2017, Cheng *et al*., 2018, Zhang *et al*., 2018). Comparison between mRNA and protein expression profiles can help uncover possible post-transcriptional (Cheng and Chen, 2004, Liu and Chen, 2016, Kawa and Testerink, 2017) and/or translational controls (Skelly *et al*., 2016, Sablok *et al*., 2017). Furthermore, the function of a given gene can be inferred by its temporal-spatial patterns associated with a given phenotype. When equipped with various high-throughput technologies, such analyses can be expanded to the whole genomic level (Brautigam and Gowik, 2010, Reinders and Paszkowski, 2010, Jackson *et al*., 2011, Matros *et al*., 2011, Schmitz and Zhang, 2011, Schneeberger and Weigel, 2011, Huang and Han, 2014). With the development of numerous bioinformatics tools, many studies have begun to employ Multi-Omics approaches in order to elucidate regulatory networks governing gene expression that consequently regulates diverse phenotypes or biological processes (Zheng *et al*., 2017, Choi, 2018, Liu *et al*., 2018, Shang and Huang, 2018).

It is notable that many genes exhibit rapid and sensitive responses to the surrounding environment, which makes collecting experimental materials from the same sample a necessity to ensure the maximum accuracy for subsequent analyses. Simultaneously extracting DNA, RNA, and proteins is possible through a guanidine isothiocyanate-based protocol (Coombs *et al*., 1990), and several derivatives using Tri Reagent (containing guanidine isothiocyanate as a major component) have been developed with minor modifications (Tolosa *et al*., 2007, Radpour *et al*., 2009, Xiong *et al*., 2011, Rajput *et al*., 2012, Pena-Llopis and Brugarolas, 2013, Vorreiter *et al*., 2016). Generally, Tri Reagent separates DNA, RNA, and proteins to different fractions. Sequential precipitation or its combination with specific nucleic acid binding columns facilitate further purification. However, almost all those protocols require protein precipitation and dissolving steps, increasing difficulties in experimental handling and reducing yields. In addition, those protocols may not be as effective for valuable samples in small quantity, which further restrains their applications in research.

Here, we established a simple and robust method to co-purify DNA, RNA, and proteins from the same sample. This method does not employ Tri Reagent, has minimum precipitation steps, and generates ready-to-use RNA and protein solutions with high quality. Our analysis showed that both nuclear and cytosolic proteins can be effectively purified. Of note, this method is compatible with samples in small quantity.

We attempted to employ the powerful *Arabidopsis* genetic resources to facilitate studies on viroid biogenesis, using the protoplast replication system. Viroids are circular non-coding RNAs that mainly infect crop plants (Ding, 2009, Flores *et al*., 2014, Flores *et al*., 2016). The host machinery and the underlying mechanisms for viroid biogenesis are largely unclear due to technical challenges in biochemical approaches. Viroids cannot embark systemic infection in *A. thaliana* (Ding, 2009), which hindered the usage of *Arabidopsis* genetic resources for viroid research. However, multiple viroids can replicate in *Arabidopsis* transgenic lines expressing their cDNAs, indicating the presence of conserved machinery for viroid biogenesis (Daros and Flores, 2004). Here, we demonstrated the replication of potato spindle tuber viroid (PSTVd) in *Arabidopsis* protoplasts and successfully applied our method to co-purify DNA, RNA, and proteins. This experimental platform will significantly enhance our capacity to probe the host machinery and the functional mechanisms for PSTVd nuclear import/export and propagation. Moreover, our new method should have broad applications for various research in plant biology and other disciplines in molecular biology.

## Results

### Recovering proteins after RNA enrichment

We chose transgenic *Nicotiana benthamiana* 16C plant with constitutive GFP expression as the test materials and used GeneJET plant RNA purification kit and MagJET RNA purification kit for RNA purification. The total RNA was purified following manufacturer’s instructions. It is notable that the MagJET family has a variety of products to purify different RNA species, which is ideal for one-step enrichments of desired RNA populations for downstream analyses. As expected, we could easily detect GFP mRNA using total RNA purified from either the GeneJET plant RNA purification kit or the MagJET RNA kit by Reverse Transcription (RT)-PCR (Figure 1).

**Figure 1.**
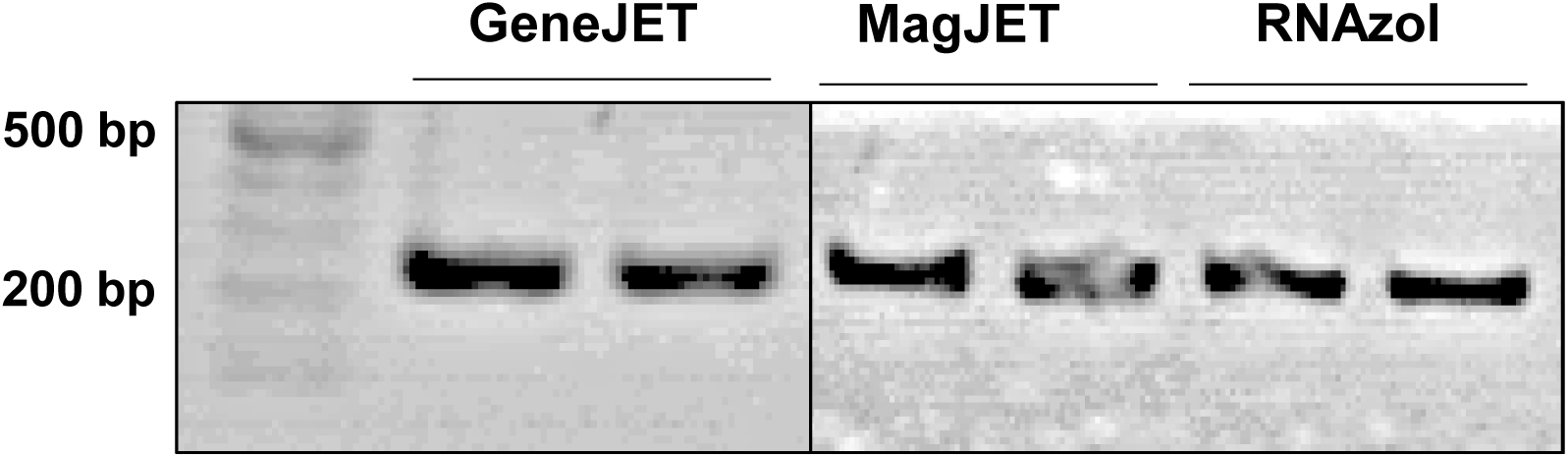
RT-PCR assessing RNA quality. RT-PCR cloning of a *GFP* fragment confirms the success purification of RNA.

Because proteins normally pass through the column or remain in the supernatant after RNA binding to the magnetic beads, we reasoned that it is possible to recover proteins from the flow-through fraction (FL) after RNA binding to the GeneJET column or the unbound supernatant fraction (Su) after RNA binding to the MagJET beads. Interestingly, the SDS-PAGE Sample Prep Kit employs DMSO-denaturing-based principle to enrich proteins and remove undesired chemicals from protein solutions. Therefore, we decided to test whether a combinational usage of both RNA and protein purification kits can purify RNA, and proteins from the same sample. We added DMSO to the FL and Su fractions and applied the mixtures to the SDS-PAGE Sample Prep Kit to enrich total proteins. After applying the enriched proteins to SDS-PAGE gel electrophoresis and silver staining, we observed effective recovery of total proteins from both the FL and Su fractions. When comparing the recovering efficiency with proteins directly purified from leaf samples using RIPA buffer, recovery from FL and Su are less than the direct RIPA purification (Figure 2). Nevertheless, most of the protein bands in RIPA buffer purified samples are present in samples prepared with our method. Furthermore, the consistent protein patterns in the replicates using our method indicates that this new method is reliable for protein analyses. As a further test, we performed an immunoblotting assay and detected the presence of GFP protein in recovered proteins using our purification method (Figure 3). Interestingly, the GFP signals were stronger in protein recovered from the FL fraction as compared with proteins directly purified using RIPA buffer, likely due to the removal of undesired chemicals by the SDS-PAGE Sample Prep Kit. Since the Su fraction already contains a high concentration of ethanol that can denature proteins, we tested if proteins can be effectively recovered using the SDS-PAGE Sample Prep Kit without supplementing DMSO. As shown in Figure 4, silver staining of total proteins and immunoblotting detection of GFP demonstrated that omitting DMSO slightly reduced the protein recovery yield but still provided a desired recovery of proteins. Since supplementing DMSO to the RNA-depleted solution significantly increases the solution volume and centrifugation steps, it is possible to omit DMSO to improve the speed of purification process for high-throughput analyses.

**Figure 2.**
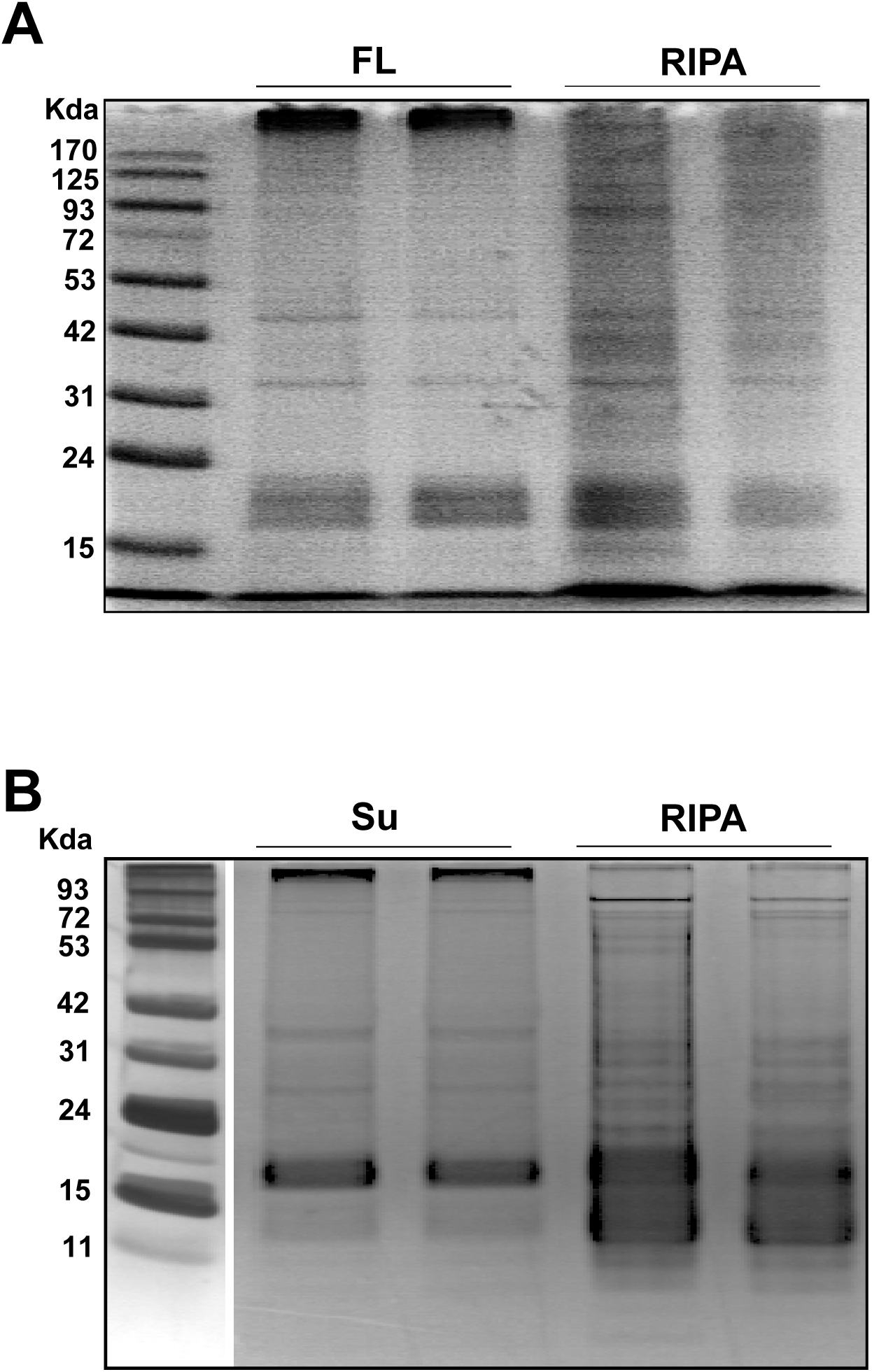
Silver staining assessing the quality of enriched total proteins. (A) the comparison of proteins enriched from the flow-though (FL) using the GeneJET RNA purification kit with direct protein purification using RIPA buffer. (B) the comparison of proteins enriched from the RNA-depleted supernatant (Su) using the MagJET RNA purification kit with direct protein purification using RIPA buffer.

**Figure 3.**
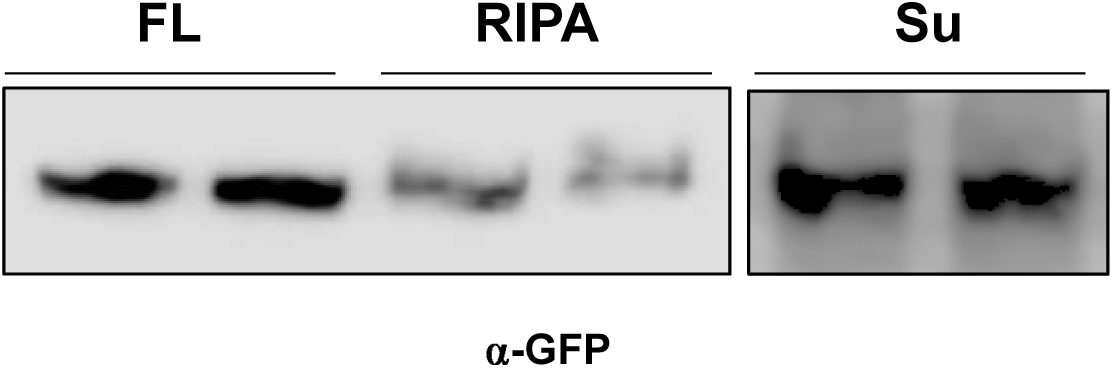
Immunoblotting detection of GFP. Immunoblotting successfully detected the GFP from our new protocol and from RIPA buffer purification. It is noticeable that our protocol enriched GFP from the flow-though of RNA purification column, supported by the better signals as compared with the RIPA purified samples.

**Figure 4.**
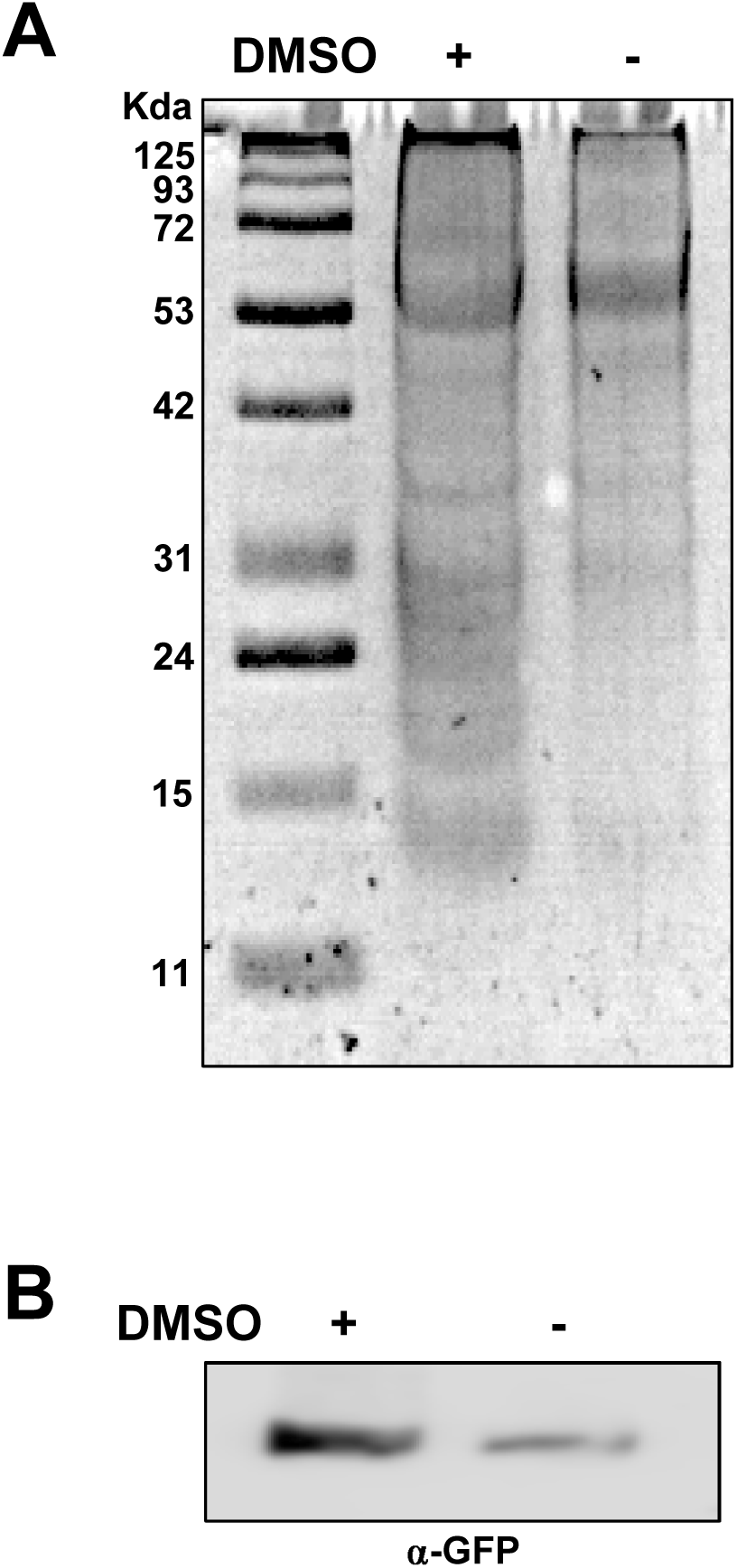
Assessing the effect of DMSO. Two equal volume of RNA-depleted supernatant from MagJET RNA purification were treated with or without DMSO. Silver staining of SDS-PAGE gel (A) and immunoblotting (B) demonstrated that the supplementation of DMSO slightly enhanced the protein recovery rate, but may be omitted to simplify the protocol for high-throughput assays.

### DNA recovery from tissue debris using DNAzol-ES

Some studies require analysis of genomic DNA. Therefore, it is desirable to purify genomic DNA from the same sample together with the purification of RNA, and proteins. We reasoned that the tissue debris as the leftover from RNA purification steps contains a sufficient amount of genomic DNA that can be purified and analyzed. We applied DNAzol-ES to the debris and followed the purification protocol from the manufacturer. From six *N. benthamiana* seedlings (<1 mg), we recovered genomic DNA from the debris (157.5 ng and 192.5 ng in two replicates). The recovery rate is about 5- to 7-fold less as compared with the direct DNAzol-ES purification (850 ng and 1090 ng in two replicates) using the same amount of starting materials. Nonetheless, it still provides a reasonable amount of genomic DNA for subsequent analyses. PCR amplification of a GFP fragment showed that the genomic DNA purified from the debris are suitable for downstream molecular analysis. It is notable that if only genomic DNA and RNA are desired for analyses, a combinational use of RNAzol and DNAzol-ES as instructed in manuals will maximize the recovery of genomic DNA and RNA.

With the successful recovery of genomic DNA from the same sample, we established an effective method to co-purify genomic DNA, RNA, and proteins from the same sample. As illustrated in Figure 5, samples are subjected to RNA purification by various commercial kits, and the flow-through fraction or the unbound supernatant are used for protein recovery using the SDS-PAGE Sample Prep Kit. Genomic DNA is recovered from tissue debris using DNAzol-ES.

**Figure 5.**
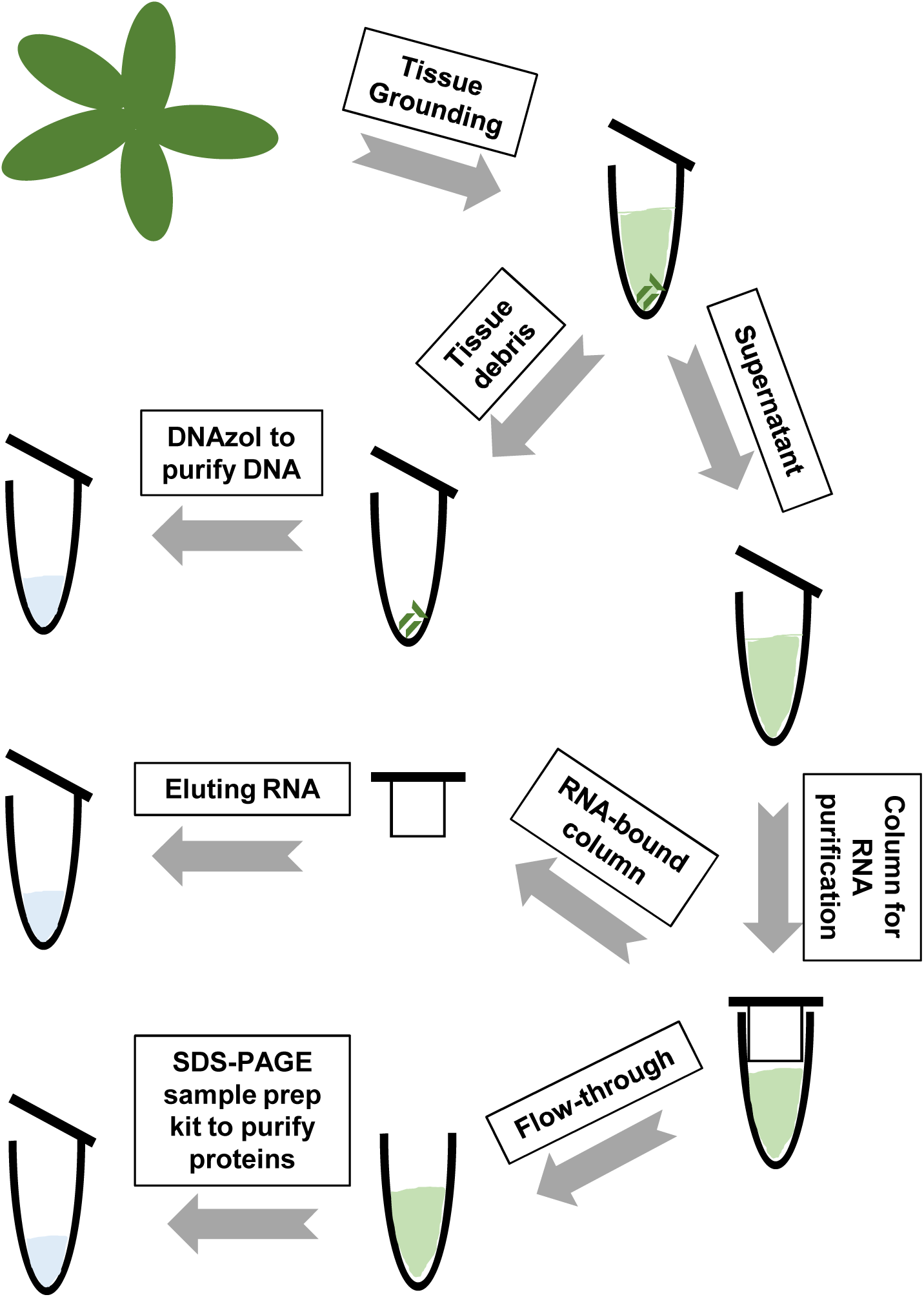
A flow chart demonstrating the procedures to co-purify DNA, RNA, and proteins.

### Co-purification of DNA, RNA, and proteins from protoplasts

We are interested in testing whether this method is efficient for samples in small quantity, such as protoplasts. Protoplast system allows rapid analysis of protein subcellular localization (Priyadarshani *et al*., 2018, Rolland, 2018), gene functions (Hamel *et al*., 2011, Li *et al*., 2013), plant responses to stresses (Asai *et al*., 2002, He *et al*., 2007), viral replication processes (Qi and Ding, 2002, Owen *et al*., 2016), etc. However, previous studies were restrained to microscopic analysis or to only analyze RNA or proteins due to the limited amount of materials. Here, we used PSTVd to co-transfect *A. thaliana* protoplasts with GFP reporter plasmids (35S::GFP). PSTVd replicates in *A. thaliana* (Daros and Flores, 2004, Itaya *et al*., 2007) but cannot achieve systemic trafficking (Daros and Flores, 2004). In our test, about 2×10^5^ protoplast cells were harvested two days posttransfection and were subjected to nucleic acids and protein purification. As shown in Figure 6a, we could detect PSTVd replication using RNA gel blots, as indicated by the presence of circular genomic RNA. Immunoblots detected the expression of GFP, with Histone H3 as a loading control (Figure 6b). Genomic PCR was successfully performed to amplify a fragment of the endogenous Transcription Factor IIIA (TFIIIA) gene (Figure 6c). These results demonstrated that this new method is efficient for materials in small quantity. Importantly, these results also showed that our new method can effectively recover proteins from both cytosolic and nuclear compartments.

**Figure 6.**
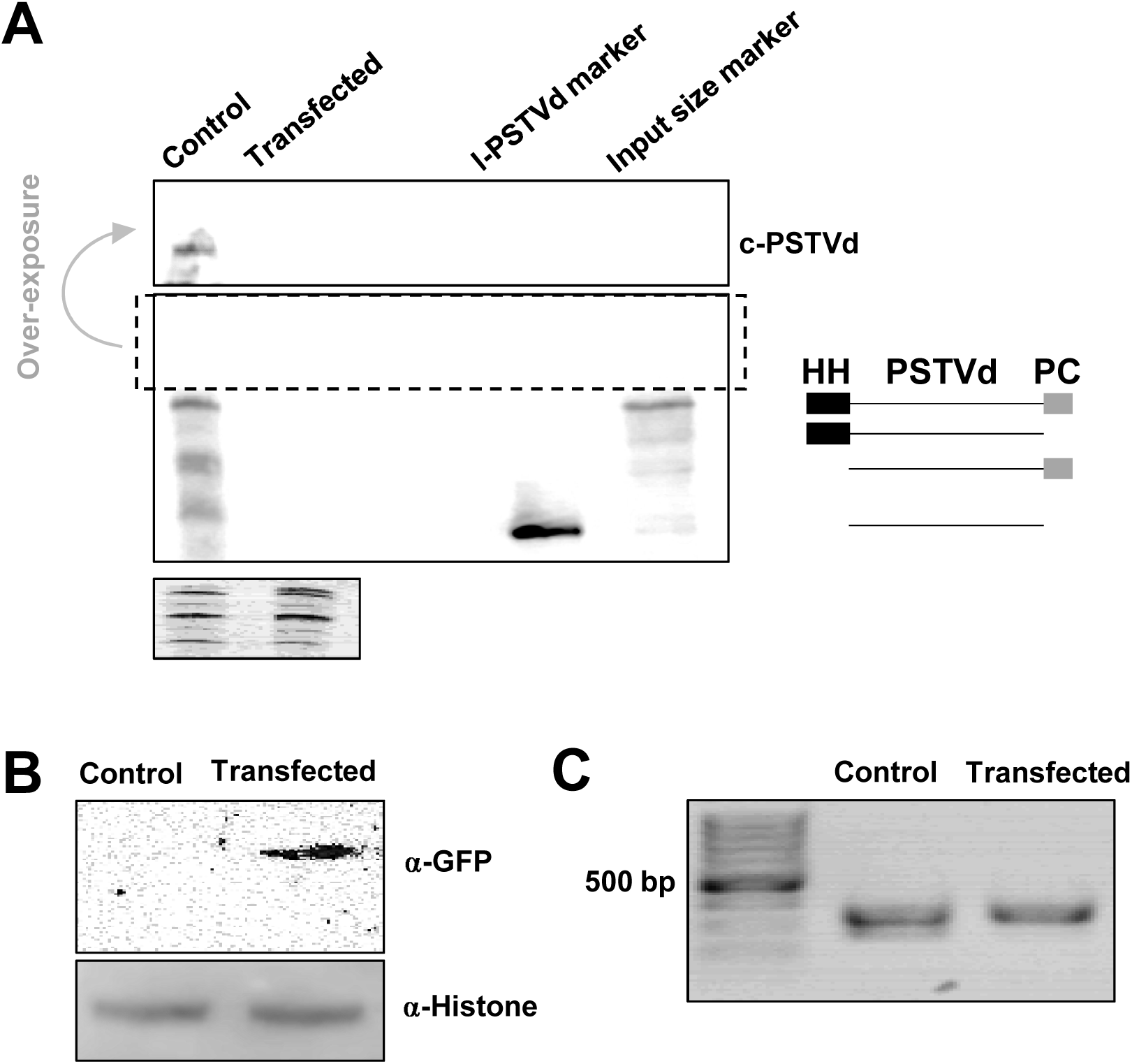
Co-purification of DNA, RNA, and proteins from protoplasts. PSTVd and 35S/.GFP co-transfected *Arabidopsis* protoplasts were harvested two days posttransfection. DNA, RNA, and proteins were co-purified using our new method. (A) RNA gel blotting detected the replication of PSTVd in *Arabidopsis* protoplasts, and the ethidium bromide staining of ribosomal RNAs served as a loading control. The size illustration for RZ-Int RNA input is on the right panel. HH and PC, hammerhead and paperclip ribozymes. (B) Immunoblotting detected the expressing of GFP, and histone H3 served as a loading control. (C) PCR using purified genomic DNA cloned a fragment from endogenous TFIIIA gene. c-PSTVd depicts circular PSTVd genome.

## Discussion

With the rapid development of various Omics technologies, there has been a trend to analyze Multi-Omics data coordinately to achieve a better understanding of the complex gene networks in various biological processes (Brautigam and Gowik, 2010). Thus, it is critical to fraction and purify distinct biological components (e.g., DNA, RNA, and protein) with high quality for analyses. It is notable that many of the genes exhibit rapid expression dynamics in response to diverse environmental stimuli, so using distinct components from the same sample will overtly enhance the accuracy of the subsequent analyses. In this regard, methods to co-purify various biological components from the same sample are necessities. Currently, most of the available protocols to co-purify DNA, RNA, and proteins from the same sample are largely based on Tri Reagent and involve multiple precipitation steps (Vorreiter *et al*., 2016). These protocols all require precipitation of proteins and then dissolving the pellets, which are time-consuming and technically challenging. Here, by using the SDS-PAGE sample prep kit, we developed a method that can purify proteins after RNA purification using various kits (column-based or magnetic bead-based). The genomic DNA can then be purified from tissue debris that is the leftover from the initial RNA purification steps. Of note, RNA was purified in the first step, which ensures the yield of high-quality RNA with minimum degradation while waiting. The robustness of the SDS-PAGE Sample Prep Kit in total protein recovery makes the method potentially compatible with a wide range of RNA purification products. This method is simple and straightforward and generates ready-to-use materials for downstream immunoblotting analysis, RNA gel electrophoresis analyses, regular cloning, or Next-generation Deep Sequencing analyses.

One advantage of our method is that it is compatible with samples in small quantities as starting materials, such as protoplasts. Protoplast assays have been widely used in plant biology to rapidly and efficiently test: 1) the cellular localization of proteins and RNAs (Priyadarshani *et al*., 2018, Rolland, 2018), 2) plant responses to biotic stresses at transcriptional and post-transcriptional levels (Asai *et al*., 2002, He *et al*., 2007), 3) protein-protein interactions (Schweiger and Schwenkert, 2014, Priyadarshani *et al*., 2018), 4) gene functions (Hamel *et al*., 2011, Li *et al*., 2013), 5) viral replications (Qi and Ding, 2002, Owen *et al*., 2016), etc. Due to the limit of sample quantity, current studies are often limited to microscopic analysis or only analyze one type of biological components (DNA, RNA, or protein) in protoplasts, undermining the value of this transient transgenic approach.

Here, we demonstrated that the combination of the *Arabidopsis* protoplast replication system and the co-purification system can significantly advance viroid research by opening the door to the powerful *Arabidopsis* genetic resources (Li *et al*., 2014). Viroids are circular noncoding RNAs that infect crop plants, often leading to plant disease (Ding, 2009, Flores *et al*., 2014). As infectious noncoding RNAs, viroids have been a productive model to dissect plant defense mechanisms against invasive RNAs (Itaya *et al*., 2007, St-Pierre *et al*., 2009, Zheng *et al*., 2017), the role of RNA three-dimensional motifs in regulating RNA systemic trafficking (Ding, 2009, Wang *et al*., 2018), and RNA-templated RNA replication by DNA-dependent RNA polymerase II (Pol II) (Rackwitz *et al*., 1981, Wang *et al*., 2016). Recent progress has begun to elucidate how viroids co-opt cellular factors to effectively propagate themselves (Flores *et al*., 2005, Ding, 2009, Nohales *et al*., 2012a, Nohales *et al*., 2012b, Minoia *et al*., 2014, Katsarou *et al*., 2016, Wang *et al*., 2016, Dissanayaka Mudiyanselage *et al*., 2018, Jiang *et al*., 2018). PSTVd, and viroids in family *Pospiviroidae*, employ Pol II for transcription (Rackwitz *et al*., 1981). A splicing variant of TFIIIA (TFIIIA-7ZF) facilitates Pol II-catalyzed transcription using PSTVd RNA template (Wang *et al*., 2016). Interestingly, the splicing of *TFIIIA* transcripts is regulated through a direct interaction between PSTVd and ribosomal protein L5 (RPL5), which results in optimal expression of TFIIIA-7ZF favoring PSTVd transcription (Jiang *et al*., 2018). During replication, the ligation of linear unit-length of PSTVd is catalyzed by host DNA ligase 1 (Nohales *et al*., 2012a). It is interesting that a PSTVd relative, hop stunt viroid, can directly interact with host Histone deacetylase 6 and alter the DNA methylation patterns in host genome (Castellano *et al*., 2016). Viroids in family *Avsunviroidae* utilize a nuclear-encoded chloroplastic DNA-dependent RNA polymerase for transcription (Navarro *et al*., 2000). During processing, a chloroplastic RNA binding protein PARBP33, binds Avocado sunblotch viroid (ASBVd) and facilitates ASBVd self-cleavage (Daros and Flores, 2002). The monomeric linear ASBVd RNA is circularized by a chloroplastic isoform of tRNA ligase (Nohales *et al*., 2012b). In spite of these progresses, future studies are required to unravel other cellular factors involved in viroid biogenesis and their functional mechanisms. Using the *Arabidopsis* protoplast system in combination with our co-purification method, viroid research can now take advantage of the potent genetic resource for future explorations. Our method present here is also applicable to various other research in molecular biology.

## Experimental procedures

### Plant growth and protoplast

*N. benthamiana* plants were grown in a growth chamber at 25°C and with a 16/8 hr light/dark cycle. *A. thaliana* plants were grown in a growth chamber at 23°C and with a 16/8 hr light/dark cycle. Using 4-week old *Arabidopsis* plants, protoplasts were isolated following a published protocol (Wu *et al*., 2009). Briefly, we used 3% cellulose (Onozuka Yakult Pharmaceutical IND., Tokyo, Japan) and 0.8% macerase (MilliporeSigma, Burlington, MA) in digestion buffer (0.4 M mannitol, 20 mM KCl, 20 mM MES, 10 mM CaCl2, 0.1% BSA, 5 mM β-mercaptoethanol, pH 5.7) to digest leaves with the lower epidermis layer removed by tapes. After 1 hr digestion, the protoplasts were pelleted using 1 min centrifugation at 150 g. The pelleted protoplasts were then sequentially incubated in the W5 buffer (5 mM MES, 154 mM NaCl, 125 mM CaCl_2_, 5 mM KCl, pH 5.7) on ice and MMg solution (4 mM MES, 0.4 M mannitol, 15 mM MgCl_2_, pH 5.7) at room temperature. About 10^5^ cells in 200 μl MMg buffer were supplemented with 40 μg 35S/.GFP plasmid and 5 μg RZ-PSTVd^Int^ RNA (Itaya *et al*., 2007), then mixed with 200 μl PEG solution (4 g PEG4000, 3.5 ml ddH_2_O, 2 ml 1M mannitol, 1 ml 1M CaCl_2_) for 5 min incubation. Finally, protoplasts were washed with W5 solution and then incubated in WI solution (0.4 M mannitol, 20 mM KCl, 20 mM MES, pH 5.7) for 2 days before RNA purification.

### RNA extraction and analysis

Leaf samples were collected in 1.5 ml microcentrifuge tubes and ground in liquid nitrogen. The GeneJET plant RNA purification kit and the MagJET RNA Kit (Thermo Fisher Scientific, Waltham, MA) were used for RNA purification as instructed in the manuals. Briefly, when using the GeneJET plant RNA purification kit, tissue lysate was loaded into the column for centrifugation. The flow-through (FL) was collected for subsequent protein purification. The column was washed twice using washing buffer from the kit. Nuclease-free water was used to elute the total RNA from the column after the DNase I treatment. When using the MagJET kit, 40 μl of MagJET magnetic beads and 400 μl of ethanol (96-100%) were added to 400 μl tissue lysate free of tissue debris, and the mixture was kept rotating for 5 min in room temperature. The mixture was then placed in the magnetic rack for 1 min. The unbound supernatant fractions (Su) were collected for protein purification. The magnetic beads were subjected to DNase I treatment and sequential washes for three times. Finally, 100μl of nuclease-free water was added to beads to elute total RNA.

The flow-through solution after RNA binding to GeneJET column and the supernatant solution collected after RNA binding to the MagJET beads were kept on ice for protein purification. The pellets tissue debris from both purification kits were saved for genomic DNA extraction. For RNA purification from protoplasts, the cells were pelleted through 1,000 g centrifugation for 2 min and then directly subjected to RNA purification using the MagJET RNA kit.

### RNA analysis using RT-PCR and RNA gel blots

For RNA analysis, total RNA extracted from above methods was subjected to reverse transcription (RT) using Superscript III reverse transcriptase (Thermo Fisher Scientific). We followed the manufacturer manual for first strand synthesis. The first strand product was subjected to PCR using primers specific for GFP mRNA (16C f: 5’-ctcccacaacgtatacatcatggc-3’ and 16C r: 5’-ccatgccatgtgtaatcccagcag-3’). For RNA gel blotting, we followed the protocol described previously. Briefly, total RNA was electrophoresed on 5% (w/v) polyacrylamide/8M urea gels for 1 hr at 200 V, transferred to Hybond-XL nylon membranes (GE Healthcare Life Sciences, Pittsburgh, PA) using a semi-dry transfer unit (Bio-Rad, Hercules, CA), and immobilized by UV cross-linking. The membrane was blocked using Denhardt solution (VWR, Radnor, PA) followed by overnight hybridization with DIG-labeled riboprobe at 65°C. DIG-labeled PSTVd specific probe was generated using HindIII-linearized pInt(-) plasmid (Qi and Ding, 2002) and a T7 polymerase MAXIscript kit (Thermo Fisher Scientific). Following the instructions of a DIG northern starter kit (MilliporeSigma), the membranes were washed and incubated with the antibody against DIG labeling. The alkaline phosphatase substrates were applied to the membranes, followed by fluorescence signal detection using C-Digit (Li-COR Biosciences, Lincoln, NE).

### Protein extraction

The FL fraction from the GeneJET kit and the Su fraction from the MagJET kit were subjected for protein purification, using the Pierce SDS-PAGE Sample Prep Kit (Thermo Fisher Scientific). Briefly, the liquid fractions supplemented with DMSO were directly loaded into a spin column. The samples were subjected to centrifugation at 2,000 g for 2 min. The flow-through was discarded. Columns were washed twice using the wash buffer and incubated with the elution buffer at 60°C for 5 min, followed by centrifugation at 2,000 g for 2 min to collect protein solutions. For RIPA buffer purification, leaf powders were mixed with 1X RIPA directly and the supernatants were used for analyses.

### Silver staining and Immunoblotting

The purified proteins were subjected to SDS-PAGE separation and transferred to nitrocellulose membranes using a semidry transfer unit (Bio-Rad). After 10 min incubation with Rapidblock solution (VWR), primary antibodies against GFP or Histone H3 (Genscript, Piscataway, NJ) were used at 1:2,000 dilution for overnight incubation at 4°C. After three washes with 1X TBST, HRP-conjugated secondary antibody against rabbit (MilliporeSigma) was added at 1:3,000 dilution for detecting Histone H3, while HRP-conjugated secondary antibody against mouse (MilliporeSigma) was added at 1:8,000 dilution for detecting GFP. After 1X TBST wash for three times and then incubation with HRP substrates (Li-COR Biosciences), the signals were captured with C-Digit (Li-COR Biosciences). For silver staining, the gels after electrophoresis were treated with the Silver Bullit^™^ kit (VWR), following instructions in the user manual.

### DNA extraction

The pellet of cell debris left from RNA purification was subject to 1 ml DNAzol-ES following instructions from the vendor (Molecular research center Inc., Cincinnati, OH). The mixture was subject to centrifugation at 10,000 g for 10 min, and the supernatant was then transferred to a new centrifuge tube. Then 500 μl 100% ethanol was added to the supernatant and mixed at room temperature for 3 min. Genomic DNA was precipitated at 5,000 g centrifugation for 5 min. The DNA pellet was washed twice with 1 ml ice cold 75% ethanol and air dried for 5 min. The pellet was dissolved using 20 μl TE buffer. The purified genomic DNA samples, after dilution to equal concentration (~10 ng/μl), were subjected to PCR analysis to detect GFP gene in 16C plants (primers described above) or TFIIIA fragment in *Arabidopsis* (AtTFIIIA genome p f: 5’-ggagacctcctgagaagctccagc-3’ and AtTFIIIA genome p r: 5’-gtccttatcacggttgtcattactatg-3’). PCR product was confirmed using agarose gel electrophoresis.

## Acknowledgements

We are grateful for David Baulcombe at University of Cambridge for sharing the *N. benthamiana* 16C line as a gift. We are thankful for Donna Gordon at Mississippi State University for constructive discussions. We thank Shachinthaka Dissanayaka Mudiyanselage at Mississippi State University for critical reading. This work was supported by US National Science Foundation (IOS-1564366) to YW; The Strategic Research Initiative fund from College of Arts and Sciences to YW; and American Heart Association under grant to BL.

